# Loss of the Central Region Reshapes the Dynamic Landscape of the Cellular Prion Protein and Its Plasma Membrane Interaction

**DOI:** 10.64898/2026.05.15.725371

**Authors:** Marta Rigoli, Pietro Faccioli, Emiliano Biasini

**Author notes:** Correspondence should be addressed to Marta Rigoli or Emiliano Biasini.

## Abstract

Prion diseases are fatal neurodegenerative disorders driven by the conversion of the cellular prion protein (PrP) into a misfolded, pathogenic conformer. Beyond serving as a substrate for prion propagation, PrP is also thought to mediate neurotoxic signaling. Within this framework, the central region of PrP has emerged as a critical regulatory element. Notably, deletion of residues 105–125 (ΔCR) leads to spontaneous neurodegeneration in vivo and induces abnormal ionic currents in cultured cells and primary neurons, indicating that this region is essential for controlling the toxicity of the N-terminal domain. Current models propose that the N-terminus functions as a toxic effector whose activity is modulated by the C-terminal domain. This intramolecular interplay is likely central to the physiological role of PrP, and its disruption may contribute to neurodegeneration. Here, we investigated how deletion of the central region affects the structure and dynamics of full-length PrP. We generated membrane-bound models of full-length, diglycosylated wild-type (WT) PrP and the neurotoxic ΔCR mutant, and compared their conformational dynamics using molecular dynamics simulations. The two proteins exhibited markedly distinct behaviours. WT PrP adopted a more compact conformational ensemble of the N-terminal domain, consistent with stabilizing interactions between the flexible N-terminus and the globular C-terminal domain. In contrast, the ΔCR variant displayed more extended conformations and a substantial redistribution of intramolecular contacts, including the loss of specific interactions between the disordered N-terminal tail and the globular domain. This altered structural organization was accompanied by an increased propensity of the N-terminal domain to approach the membrane surface in the mutant. Our results provide a molecular model in which the central region engages intramolecular interaction networks that ultimately help regulate N-terminal residence at the plasma membrane, offering mechanistic insight into how CR deletion shifts the conformational ensemble toward membrane-associated states that may be associated with neurotoxic activity.

## Introduction

Prion diseases, or transmissible spongiform encephalopathies, are fatal neurodegenerative disorders caused by the conformational conversion of the cellular prion protein (PrP) into a misfolded, pathogenic isoform (PrP^Sc^) ^1^. These disorders affect humans and animals and share mechanistic parallels with other protein misfolding diseases, including Alzheimer’s and Parkinson’s diseases ^2^. Central to prion pathogenesis is the remarkable conformational plasticity of PrP, a glycosylphosphatidylinositol (GPI)-anchored membrane protein composed, in its mature form, of residues 23-230. The protein encompasses a highly flexible N-terminal domain (residues 23-125) and a structured C-terminal globular domain (residues 126-230) containing three α-helices and two short antiparallel β-strands. The N-terminal region of PrP plays a critical role in both physiological function and neurotoxicity. This segment includes the octapeptide repeat (OR) domain (residues 59-90), which binds divalent metal ions such as Cu^2+^ and Zn^2+^ via histidine coordination. Metal binding promotes a regulatory intramolecular interaction between the flexible N-terminus and a negatively charged surface on the C-terminal domain, involving residues located primarily on helices 2 and 3 ^3^. This cis interdomain interaction suppresses the intrinsic toxic activity of the N-terminal effector domain, which otherwise can induce membrane permeabilization, spontaneous ionic currents, and neuronal dysfunction ^2,4–11^. Disruption of this regulatory mechanism through mutation, altered metal coordination, or post-translational modification has been linked to enhanced neurotoxicity and familial prion disease ^9,12^. A particularly informative model of disrupted intramolecular regulation is the ΔCR (central region deletion) mutant of PrP. This mutant lacks a conserved stretch of residues within the N-terminal region (residues 105-125), a segment that lies between the OR domain and the structured C-terminus ^10^. The central region has been proposed to function as a critical regulatory element that modulates the orientation, dynamics, and membrane interactions of the N-terminal domain ^13^. Deletion of this region produces a protein that exhibits dramatically enhanced neurotoxicity in cultured cells and in vivo models, including the generation of spontaneous inward cationic currents and increased sensitivity to membrane perturbation ^7^. Importantly, the magnitude of these toxic phenotypes exceeds that of many pathogenic point mutations, underscoring the central region’s key role in maintaining PrP in a neuroprotective conformation ^8^. Despite extensive biochemical and electrophysiological characterization, the structural and dynamical consequences of central region deletion remain incompletely understood. Experimental structural approaches are challenged by the intrinsic flexibility of the N-terminal domain and by the membrane-associated, post-translationally modified nature of PrP. While biophysical studies have provided insights into interdomain contacts and surface interaction patches on the C-terminal domain, these techniques offer limited spatial and temporal resolution for capturing the full conformational ensemble sampled by PrP variants, particularly in the context of large-scale domain rearrangements or membrane-proximal conformations.

Molecular Dynamics (MD) simulations provide a powerful complementary approach for dissecting the structural basis of ΔCR-mediated toxicity. By resolving atomic-level motions over time, MD enables direct comparison of the conformational landscapes of WT and deletion mutants, characterization of transient interdomain contacts, quantification of secondary structure stability, and identification of altered electrostatic and solvent-exposed surfaces.

In this study, we use atomistic MD simulations to determine how deletion of the central region (Δ105–125) alters the conformational ensemble and intramolecular regulation of native PrP. Notably, this represents, to our knowledge, the first MD investigation of a full-length, di-glycosylated, GPI-anchored PrP embedded in a plasma membrane-like lipid bilayer. By explicitly modelling the structured C-terminus, the intrinsically disordered N-terminal tail, complex N-linked glycans, the GPI anchor, and a POPC-cholesterol membrane environment within a single atomistic framework, we capture a level of biological realism not previously explored computationally. Such an integrated, atomistic approach allows us to provide a single-residue-resolution mechanism of how the CR deletion rewires intramolecular interaction networks and biases the N-terminus toward membrane-associated conformations, events that are paradigmatic of its intrinsic neurotoxicity.

## Methods

### Model building

PrP molecular models were built by combining two complementary sources: the experimental WT PrP structure (PDB ID: 7FHQ) and the AlphaFold-predicted model AF-P04156-F1-model_v3 ^14^. The C-terminal domain, together with the adjacent portion of the N-terminal region (residues 91–230), was derived from the 7FHQ structure, whereas the remaining intrinsically disordered N-terminal segment (residues 23–90) was obtained from the AlphaFold model. The complete membrane-bound systems, comprising the protein, N-linked glycans, a GPI anchor, and a model plasma membrane, were assembled using the CHARMM-GUI web server ^15^. The membrane was composed of POPC (80%) and cholesterol (20%) and had approximate dimensions of 150 × 150 Å. The GPI anchor was linked to residue S230. Residues N181 and N197 were decorated with complex N-glycans, each one composed by four N-acetylglucosamines, three mannoses, two galactoses, one fucose, and two sialic acids. The CHARMM36/CHARMM36m force fields were used to generate the topology and parameters for all systems ^16^. The ΔCR PrP variant, in which residues 105–125 were deleted, was generated by manually modifying the WT structure. After deletion of residues 105–125, the two remaining polypeptide segments were covalently rejoined to produce the final construct.

### Simulation Protocol

Molecular Dynamics (MD) simulations were performed with GROMACS 2022.1 ^17^ on the supercomputer Finisterrae III of CESGA (Centro de Supercomputación de Galicia). Protein atoms, ions, and lipids were parameterized with the CHARMM36m/CHARMM36 force field ^16^, whereas for solvent molecules the TIP3P water model was used. The charge of each system was neutralized and further Na^+^ and Cl^−^ ions were added to reproduce a physiological salt concentration of 150 mM. Simulations were performed at 310 K under temperature control by Nosé-Hoover thermostat ^18^ with a coupling constant of 1 ps, while pressure was maintained at 1 bar through the Parrinello-Rahman barostat using a 5 ps coupling time ^19^. Periodic boundary conditions were imposed in all directions. Long range electrostatic interactions were calculated with the Particle Mesh Ewald approach ^20^, adopting a short-range cut-off of 1.2 nm and a Fourier grid spacing of 0.12 nm. A 1.2 nm cut-off was also applied to van der Waals interaction calculations, using the Lennard-Jones potential. Bonds involving hydrogen atoms were constrained using LINCS algorithm ^21^, and the leap-frog Verlet integrator was used with a 2 fs integration step ^22^.

### Molecular Dynamics simulations

After construction of the protein models, the isolated PrP structure was first relaxed by all-atom MD simulations through two independent 200 ns runs. Representative equilibrated conformations were then extracted and used as starting structures for the assembly of the complete membrane-bound systems. The complete models of WT PrP and ΔCR PrP variant, di-glycosylated, and attached to the plasma membrane, were simulated for a total of four independent MD replicas of 1 µs each, for a total of 8 µs.

### Data analysis

*Root mean square deviation* (RMSD) and distances between K23 and plasma membrane were calculated by using Gromacs 2022.1 ^17^.

*End-to-end distances* were computed from four independent replicas of the WT and ΔCR systems using Gromacs 2022.1, as the distance between the centers of mass of the N-terminal residue, K23, and the tail’s C-terminal residue of each construct that is residue 125 for WT molecule and residue 104 for ΔCR molecule.

To provide a reference for comparing the two ensembles, we also modeled each N-terminal segment as a worm-like chain (WLC), a minimal exactly solvable model for semiflexible polymers. The essential parameters of the model are the contour length, (L), and the persistence length, l_p_ ^23^.

The contour length was defined as *L* = (*n*_*res*_ − 1)*a*, where *n*_*res*_ is the number of amino acids in the segment (WT: 23–125; ΔCR: 23–104), and *a* = 0.38 nm is the typical distance between consecutive C_α_ atoms along the backbone. The persistence length controls the polymer stiffness and corresponds to the characteristic distance along the chain over which orientational correlations decay. A typical value for unstructured peptide chains is *l*_*p*_ ∼ 0.6 nm^24^.

In the WLC model, the mean-squared end-to-end distance is given by:

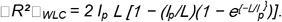

An effective persistence length (l_p-eff_) was obtained independently for both WT and ΔCR molecules by matching the WLC prediction to the simulation ensemble average. l_p-eff_ was calculated by numerically solving □*R*^*2*^ □*WLC*(L,*l*_*p*_ ) = □R^2^□sim using a bisection root-finding procedure.

*Contact map* calculations were done by means of an in-house Python script. For each trajectory, contact maps were calculated from C_α_-C_α_ distances using a cutoff of 7.5 Å. The contact maps from all frames were then averaged over time to obtain one overall contact map per trajectory.

*Inter-residue contacts* involving the N-terminal region, either within the N-terminal region itself or between the N-terminal and C-terminal regions, were quantified from the MD trajectories using a distance-based contact criterion. For the more detailed analysis of N-terminal/N-terminal and N-terminal/C-terminal contacts, a stricter C_α_ -C_α_ cutoff of 6 Å was used to focus on closer and more persistent contacts and to reduce noise from transient interactions in the disordered N-terminal region. Residue numbering was kept consistent across systems. For WT molecule, the N-terminal was defined by residues 23-125 and the C-terminal by residues 126-230. For the ΔCR molecule, residues 105-125 are deleted and the N-terminal therefore spans residues 23-104, while the C-terminal remains 126-230. Contacts between residues separated by fewer than four positions in sequence were excluded to avoid local-neighbor effects. As a sensitivity check, the same contact analysis was repeated using a 7.5 Å cutoff. As expected, this increased the absolute number of detected contacts, but the qualitative behavior remained consistent with the results obtained using the 6 Å cutoff.

For each residue pair (i, j), the contact occupancy was computed as *f*_*ij*_ *= n*_*ij*_ */ N*_*frames*,_ where *n*_*ij*_ is the number of trajectory frames in which residues *i* and *j* satisfy the contact criterion, and *N*_*frames*_ is the total number of analyzed frames.

To identify residues that act as inter-domain “hubs”, a residue-based N-terminal/C-terminal contact score was computed by summing pair occupancies over all partners in the opposite domain.

For a C-terminal residue j:

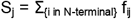

analogously for an N-terminal residue i:

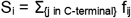

This score can exceed 1, reflecting the possibility, for a single residue, of having multiple contacts at the same time.

Three complementary visualizations were used:

i. Top N-terminal/C-terminal contact pairs, ranked by f_ij_ ;
ii. Bipartite N-terminal/C-terminal contact networks in which nodes represent residues and edges represent contacts, displayed only when f_ij_ ≥ 0.3;
iii. Top C-terminal residues ranked by the contact score S_j_.

All analyses and plotting parameters were kept consistent between WT and ΔCR to enable direct comparison.

*Secondary structures* were identified with the timeline tool in VMD ^25^ and the plots were done by means of a python script.

All the protein structure images were produced by using UCSF ChimeraX ^26^.

## Results

### MD simulations of full-length, glycosylated and membrane-anchored WT and mutant ΔCR PrP

PrP models were generated by combining an experimentally resolved structured core with an AlphaFold-predicted intrinsically disordered N-terminal region. The complete membrane-bound systems, including the protein, GPI anchor, N-linked glycans, and a lipid bilayer, were assembled using CHARMM-GUI. The membrane mimics a plasma membrane environment, and the protein was modeled with its native post-translational modifications, including glycosylation at two sites and GPI anchoring (Figure 1). The calculation of the RMSD from the initial structure of the C-terminal globular domain of the WT and ΔCR PrP along four independent 1 µs-long MD trajectories detected no major conformational changes (Figure 2). The only exception was found in a single run of ΔCR PrP, which presented a pronounced bending of helix-3. However, such a local distortion did not alter the overall folding of the protein, as shown also by the secondary structure analysis (Supplementary Figure 1), indicating no structural destabilization at the μs time scale in either WT or ΔCR PrP due to the interaction of the globular domain with glycans, membrane, or the disordered N-tail.

**Figure 1.**
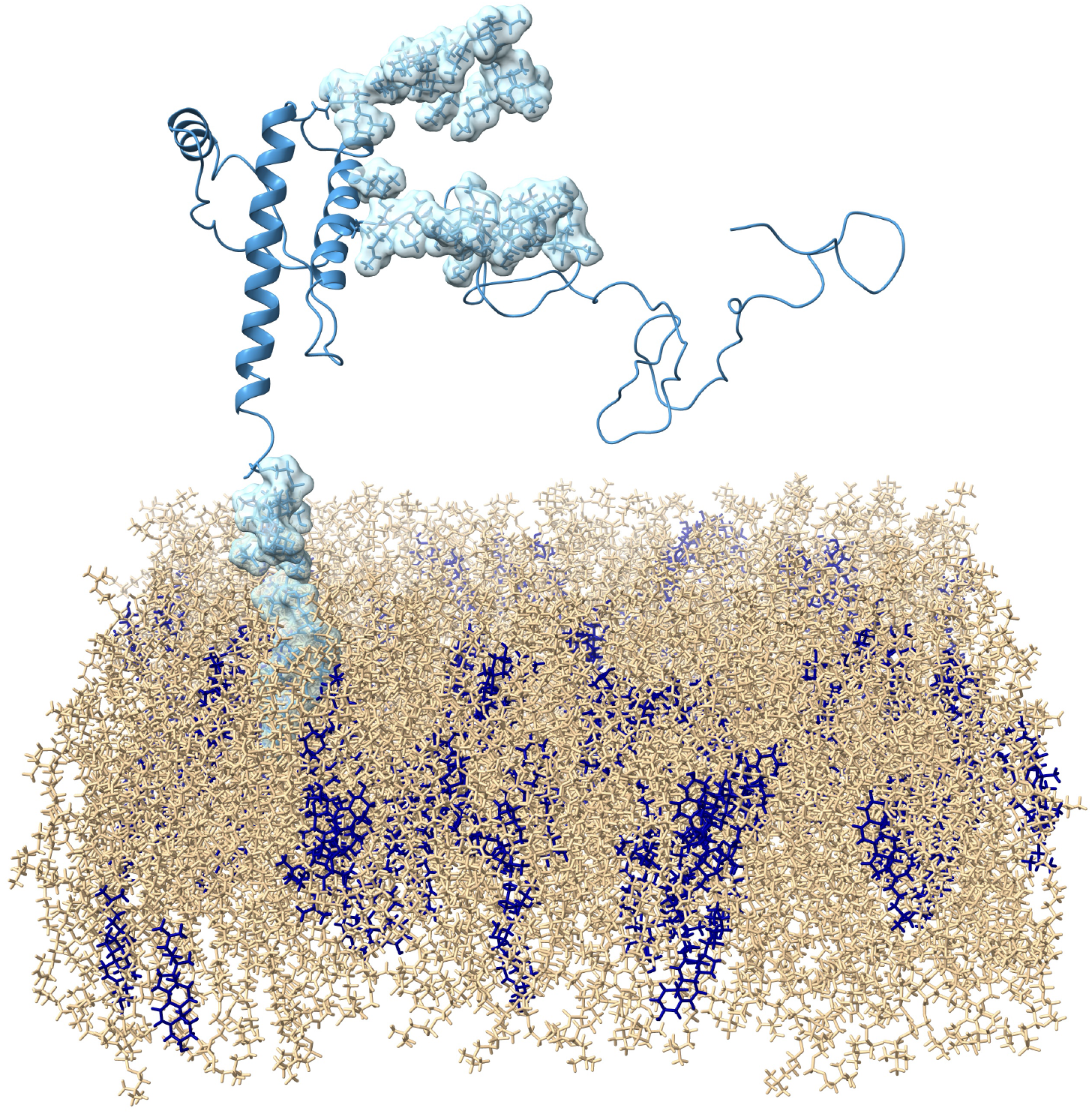
Representation of the PrP system. The PrP polypeptide is represented in blue, glycans and GPI-anchor surface are depicted in light blue, the plasma membrane is shown in tan (POPC) and in dark blue (Cholesterol).

**Figure 2.**
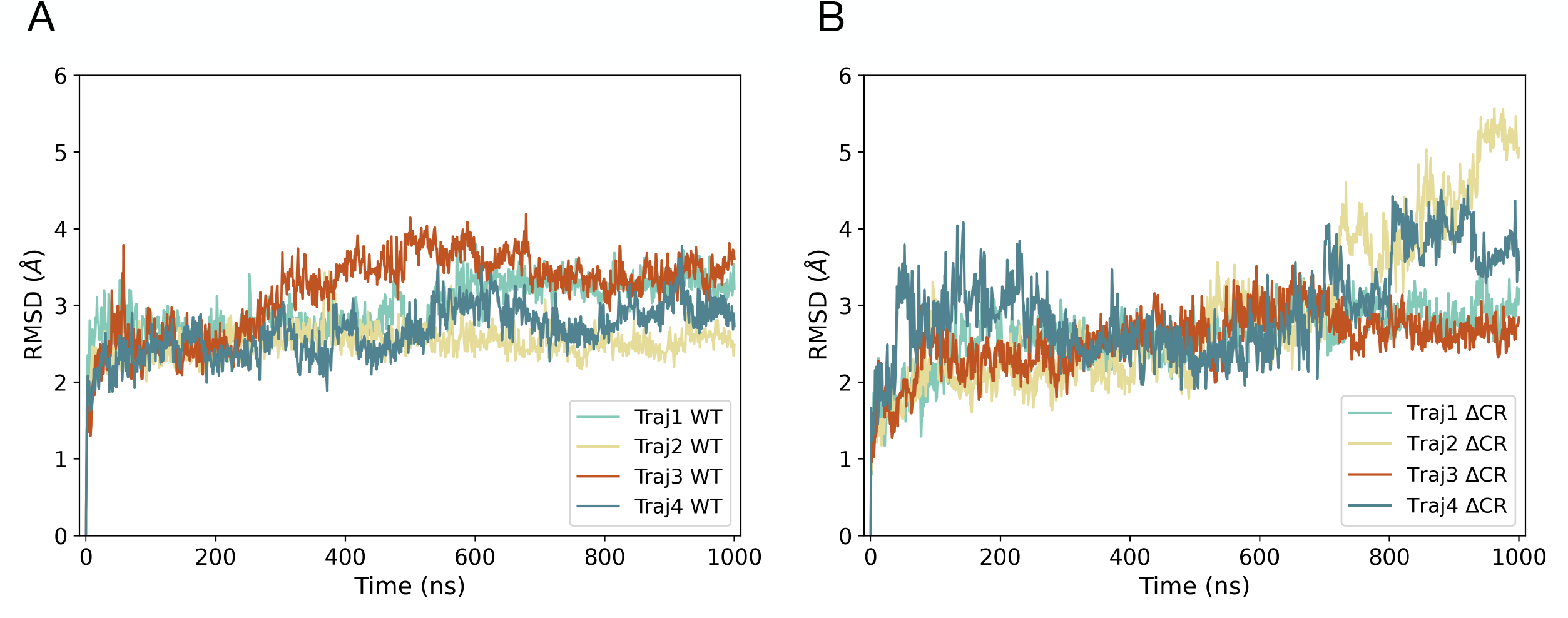
Timeline representation or the globular domain C-alpha atoms RMSD (residues 126-230). Results for WT and ΔCR PrP are reported in panels A and B, respectively. The graphs indicate a general stability of the protein structure with a deviation lower than 4 Å, with the exception of trajectory n. 2 and trajectory n. 4 of ΔCR system that present a maximum of 5.5 Å, and 4.5 Å deviation, respectively. In trajectory 2 (ΔCR) the high RMSD, observed after the timepoint around 700 ns, arises from a bending of α-helix 3 relative to the initial structure.

### The N-terminal tails of WT and mutant ΔCR PrP differ in the propensity to interact with the plasma membrane

Previous data indicate that PrP molecules carrying mutations or deletions in the central region could exert neurotoxic effects by destabilizing the integrity of the plasma membrane ^8^. In order to test this hypothesis in our system, we sought to estimate the propensity of WT or ΔCR PrP forms to interact with the outer surface of the membrane. First, we analyzed the distance of the first amino acid of the N-terminal tail (K23) of both proteins to the heads of the lipid bilayer. The distance distributions calculated for each of the four MD trajectories revealed significant differences between the WT and the ΔCR mutant. K23 exhibited longer distance from the membrane in the WT form, whereas in the ΔCR mutant, the amino acid was consistently closer. This observation indicates a greater propensity of the N-terminal region of the ΔCR mutant to approach the plasma membrane, as compared to the WT form (Figure 3). Furthermore, contact time analysis revealed a statistically significant difference in the interaction between K23 and the plasma membrane (Figure 4). Indeed, K23 of ΔCR PrP maintains prolonged contact with the outer surface of the lipid bilayer as compared to the WT protein, suggesting that the deletion of the central region alters the topology of PrP N-terminus.

**Figure 3.**
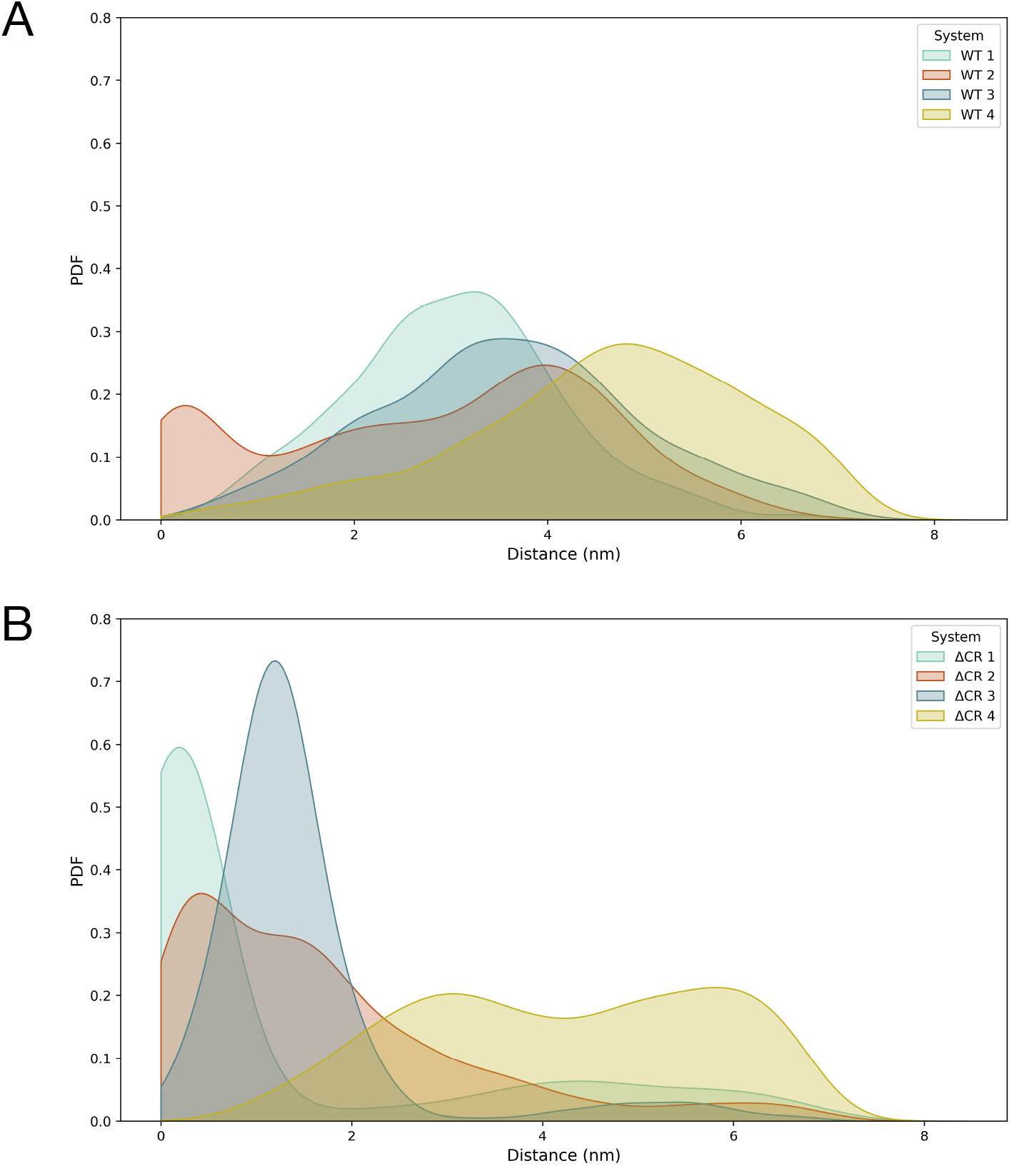
Probability density distributions of the distance between K23 and the plasma membrane for four independent WT (A) and ΔCR (B) replicas. WT PrP molecule’s trajectories are characterized by broader distributions centered at intermediate distances between 3 and 5 nm, whereas ΔCR PrP trajectories display greater heterogeneity, with some replicas showing short-distance states, below 1 nm, and others spanning a wider range of distance.

**Figure 4.**
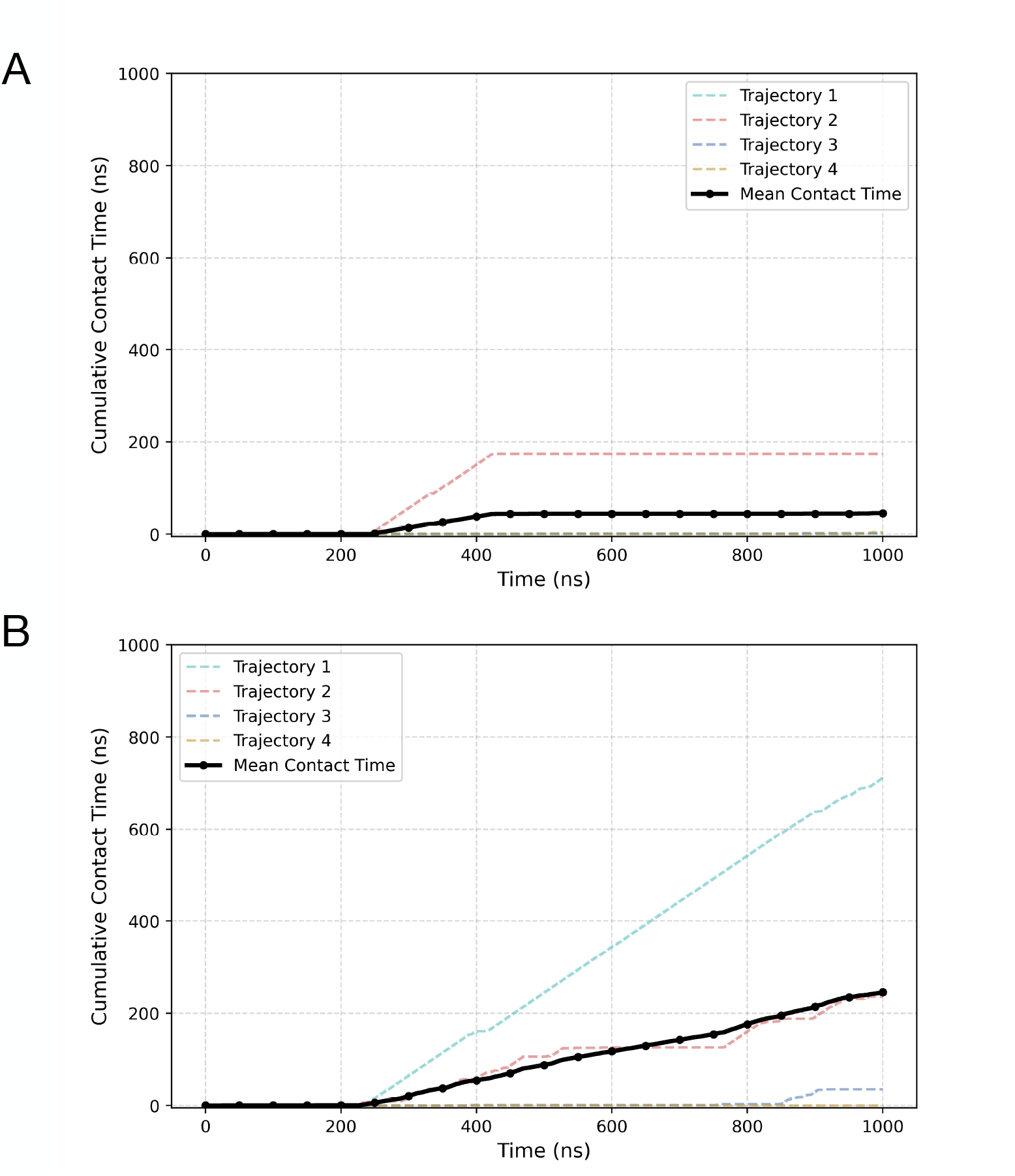
The panels show the cumulative contact time of K23 with the membrane for each trajectory as well as the mean contact time, in black. WT and ΔCR PrP molecules are shown in panels A and B, respectively. The graphs show a significant increase of the N-terminus contact time with the membrane in the ΔCR PrP molecule with respect to WT PrP.

To further corroborate these data, we analyzed the distance between K23 and the first residue of the globular domain of both proteins (G126, so called end-to-end distance). To provide a minimal polymer-physics reference for comparing the two ensembles, we modelled each N-terminal segment as a worm-like chain (WLC) of contour length L and persistence length l_p_. The contour length was estimated as L = (n_res_ − 1)a, with n_res_ being the number of residues in each segment (WT, 23–125; ΔCR, 23–104), and a = 0.38 nm as an average length per residue. The root-mean-square end-to-end distance, referred to as R_RMS_, for WLC modelled on WT or ΔCR PrP appeared as markedly shifted toward smaller values, as compared to a neutral-chain WLC baseline (l_p_ = 0.60 nm). These data indicate that the N-terminal tail of both WT and ΔCR PrP are more compact than a theoretical, random chain (Figure 5). However, despite the WT N-terminus being longer, it exhibited a smaller R_RMS_ than the ΔCR mutant (R_RMS_ for WT = 3.73 nm; R_RMS_ for ΔCR = 4.07 nm). This observation implies that differences in the effective flexibility dominate over the simple chain length. Consistent with these data, estimation of an effectiveness parameter (l_p-eff_) derived from the fitting of the WT or ΔCR PrP distributions over theoretical R_RMS_ values for a neutral-chain WLC baseline, showed lower l_p_,_eff_ for WT than for ΔCR (l_p_,_eff_ WT ≈ 0.18 nm ; l_p_,_eff_ ΔCR ≈ 0.27 nm). Collectively, these data indicate that the CR deletion reshapes N-terminal conformational dynamics, promoting a more extended and lipid-associated state consistent with its proposed membrane-disrupting activity.

**Figure 5.**
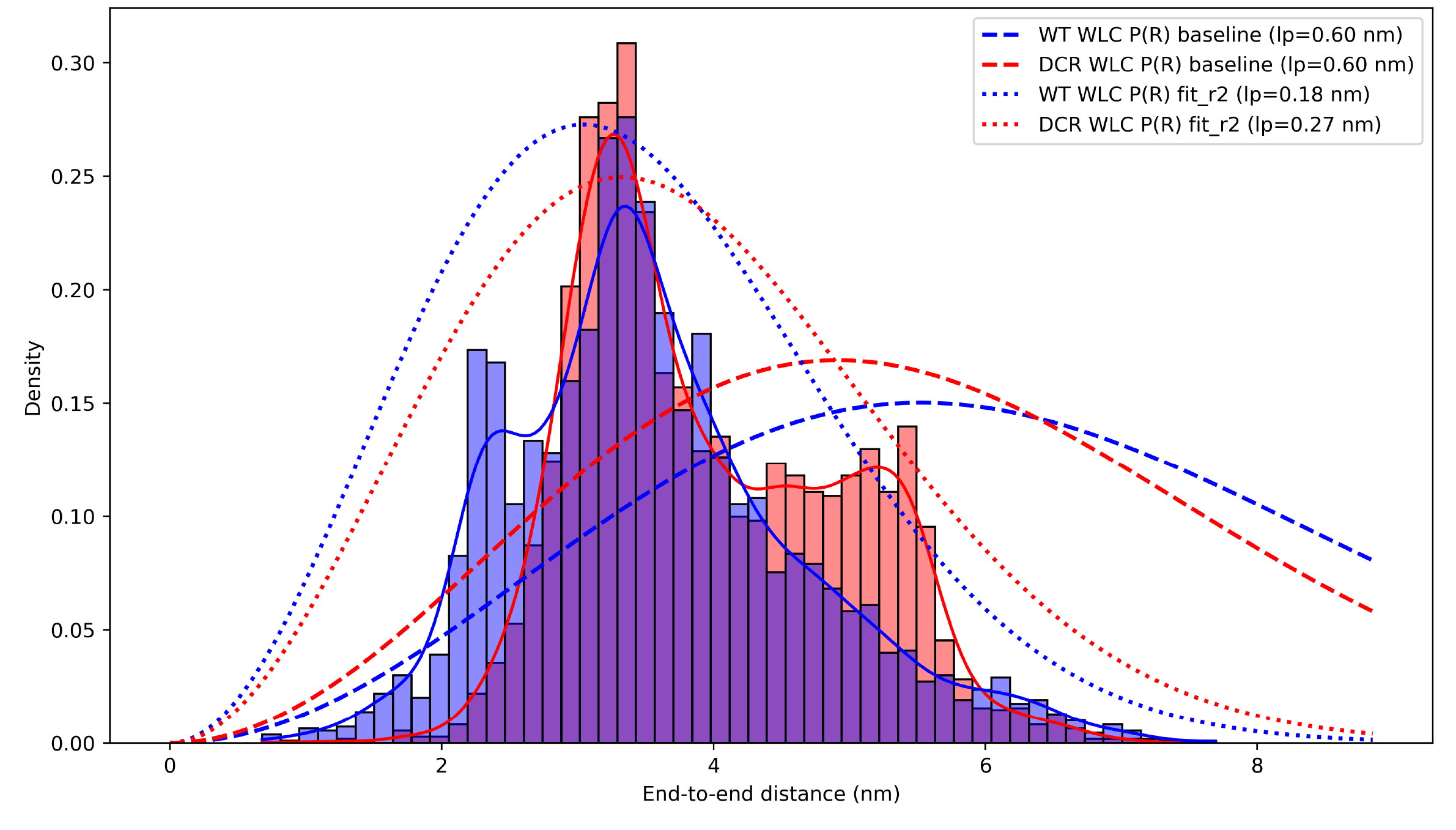
Distribution of end-to-end distances for WT (light blue) and ΔCR (light red) PrP molecules. Bars show normalized histograms over four replicas per system; solid curves show kernel density estimates. Dashed lines report the Worm-like Chain (WLC) reference distributions expected for neutral semiflexible polymers having the same contour length L as each segment (WT: residues 23–125; ΔCR: residues 23–104) using a common baseline persistence length lp = 0.60 nm. Dotted lines show WLC distributions computed using an effective persistence length lp_eff obtained by matching the WLC mean-squared end-to-end distance to the simulation, □R^2^□ _WLC_(L, l_p-eff_) = □R^2^□ _sim_, resulting in WT l_p-eff_ = 0.18 nm, ΔCR l_p-eff_ = 0.27 nm. With respect to the WLC baseline, both systems are shifted toward smaller R, indicating high compaction; ΔCR PrP is shifted to larger R than WT PrP and exhibits a broader tail, consistently with a lower structure compaction.

### WT and ΔCR PrP show different intramolecular interactions

Previous observations indicate that PrP may exhibit transient intramolecular interactions between its intrinsically disordered N-terminal domain and structured C-terminal globular domain ^27^. These contacts, largely electrostatic and modulated by copper binding, help stabilize the native conformation and may regulate susceptibility to prion misfolding. In order to test whether WT and ΔCR PrP differ in the intramolecular interactions occurring between the N-terminal flexible tail and the C-terminal globular domain, we computed contact maps. A contact map is a matrix representation that indicates which pairs of residues are found to be in contact during an MD simulation. Each contact is defined using a distance cutoff (7.5 Å for C_α_), allowing the map to summarize the average structural organization in intramolecular interactions. Contact maps computed for each trajectory of both WT and ΔCR PrP revealed differences in the interaction pattern between the N-terminal tail and the C-terminal domain of the proteins (Figure 6). Indeed, in the WT protein, persistent interactions were observed between amino acids located in the N-terminus and residues 190-200 of the globular C-terminus, belonging to helix-2 and the loop between helix-2 and helix-3. Conversely, in ΔCR PrP, the interactions involved primarily the N-terminus and the loop connecting helix-1 and helix-2, as well as the C-terminal region of helix-3. Interestingly, these data reveal that, in contrast to WT PrP, the N-terminal tail of the ΔCR mutant tends to interact with regions in the C-terminal domain lying closer to the plasma membrane. The only exception to this pattern is observed in trajectory n. 4 of the ΔCR molecule, which exhibited WT-like contacts between residues 48-50 and 189-193. Notably, in this case the cumulative persistence of K23 on the membrane was lower than the average of all the ΔCR simulations (Figure 4).

**Figure 6.**
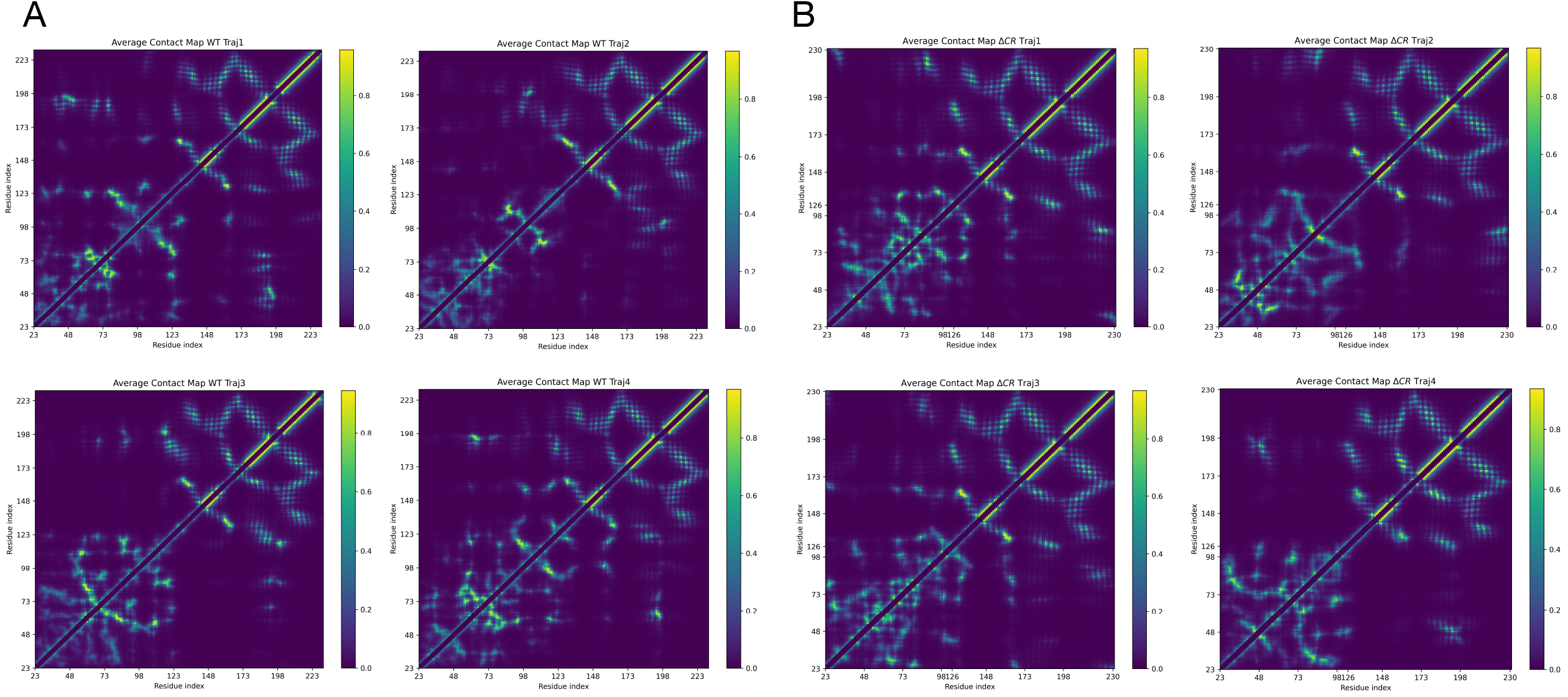
Contact map representation of the WT (A) and ΔCR PrP (B) molecules. For each trajectory an overall contact map was calculated by time averaging the contacts. The contact maps are dominated by strong near-diagonal signals, while weaker interactions are found off-diagonal. The upper-right quadrant, of each contact map, corresponds to the structured C-terminal helical region and reflects the packing of the three α-helical segments. This pattern is broadly conserved across all trajectories. In the lower-right quadrant, where contacts between C-terminal and N-terminal residues are shown, several maps of the WT PrP molecule display distinct localized hotspots that are not observed in the ΔCR PrP molecule’s maps, where the signal appears overall weaker and less structured.

To fully dissect the distinct topology of WT and ΔCR PrP, we analyzed intramolecular contacts at single–residue resolution across the trajectories. In the WT protein, the N-terminus/C-terminus interface is dominated by a high-persistence contact hotspot on the C-terminal domain centered around residues 192–200. Among C-terminal residues ranked by contact score, T193 emerges as the key contact point, followed by T199, G195, T192, and E200 (Supplementary Figure S2). This pattern indicates that the 192–200 segment of the C-terminal domain of WT PrP forms a multi-contact interaction patch rather than a single isolated contact with the N-terminal tail. In the ΔCR mutant the N-terminus/C-terminus interface is differently organized. The dominant C-terminal interaction hotspot shifts away from the 192–200 segment to residues 131–135. Here, S132 becomes the highest ranked C-terminal residue, followed by M134, S135, A133, and G131. However, a secondary interaction region emerges around residues at the extreme C-terminus, in the region 220– 228 (E221, A224, Q227, R220, and R228). Consistent with these observations, pairwise analysis obtained by building bipartite networks for WT or ΔCR showed a marked difference between the two molecules (Figure 7). The network appears immediately much denser for WT than for ΔCR PrP (pairwise contacts showing a thresholds ≥ 0.3 were 42 for WT and 23 for ΔCR PrP). For WT PrP most frequent pairs involved residues 50-51, 62-64, 101-103, and 115-117 in the N-terminus and region 192-200 in the C-terminus. Conversely, most frequent pairs in ΔCR PrP involved several individual residues spanning the most proximal region of the N-terminus and regions 131–135 and 220–228 in the C-terminus. Importantly, the analysis of intramolecular interactions occurring within the N-terminus of WT revealed a critical cluster involving residues before amino acid 100 and the CR segment (residues 105– 125; Supplementary Figure S4). The absence of the CR region in ΔCR PrP redistributes the intramolecular interactions of the N-terminus toward regions around residues 30-40 and 80-90. Overall, these data indicate that WT and ΔCR PrP adopt fundamentally different intramolecular interaction topologies. In WT PrP, the N-terminal tail forms a dense, persistent interaction network centered on residues 192–200 of the C-terminal domain and is structurally coupled to the CR segment, stabilizing a specific N–C interface. In ΔCR, removal of the CR rewires the network, with contacts shifting toward membrane-proximal C-terminal regions (131–135 and 220–228), the N–C interface becoming less dense, and N-terminal interactions redistributing internally. Therefore, the CR deletion appears to not simply weaken interactions, but instead completely reshaping the global intramolecular pairing network of PrP, resulting in a more extended and flexible N-terminal tail which results in higher tendency to interact with the plasma membrane (a video representing a single trajectory for both WT and ΔCR PrP is shown in Supplementary V1).

**Figure 7.**
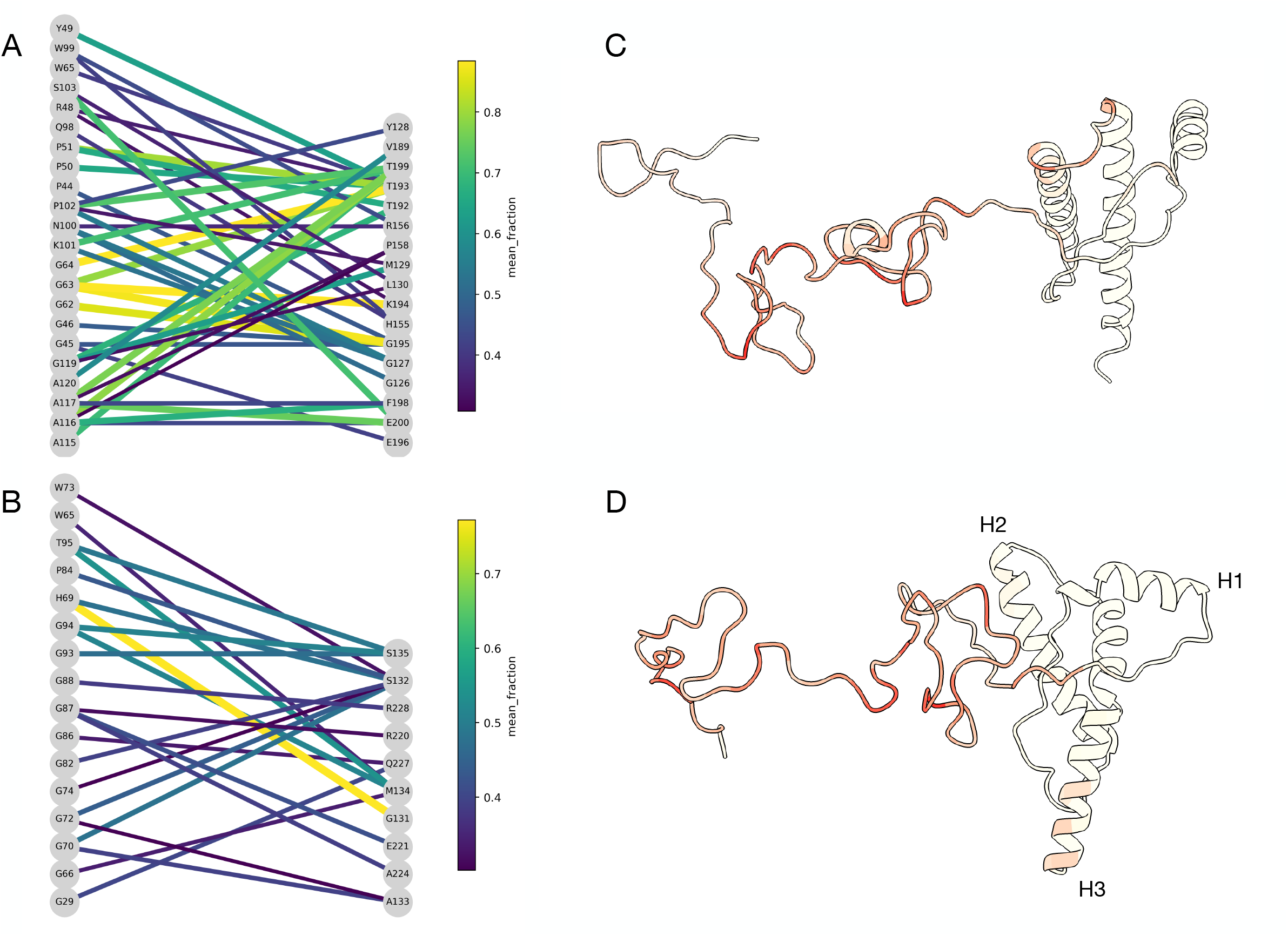
Bipartite N/C contact networks for WT and ΔCR PrP are shown in panels A and B, respectively. Nodes represent residues (N-terminus on the upper panel, C-terminus on the lower panel). Edges represent inter-domain contacts and are shown only for mean contact fraction ≥ 0.3; edge color encodes the mean contact fraction. The WT PrP network is denser and centered on the C-terminus 192–200 hotspot, while ΔCR PrP displays a sparser network with hubs relocated to 131–135 and additional connections near 220–228. Contact frequency representation for WT and ΔCR PrP are shown in panels C and D, respectively. The residues are colored by their contacts score, with darker colors indicating more persistent interactions. Contacts within the C-terminus are excluded.

## Discussion

In this work, we performed MD simulations of full-length, glycosylated, and membrane-anchored WT and ΔCR PrP to investigate the structural and dynamic effects of central region deletion. We found that the C-terminal globular domain remains stable in both forms, indicating that neither glycosylation, membrane anchoring, nor the disordered N-terminal tail induces structural destabilization. In contrast, the N-terminal region exhibits distinct behaviors: over the μs time scale which was accessible to our MD simulations, the ΔCR mutant shows an increased propensity to approach and interact with the membrane and adopts a more extended conformation compared to the compact WT. These differences are accompanied by a substantial reorganization of intramolecular interactions, with WT PrP displaying a dense N–C interaction network centered on residues 192–200, while ΔCR PrP shows a redistributed and less connected network involving membrane-proximal regions. Overall, our results indicate that deletion of the central region does not disrupt global folding but reshapes N-terminal dynamics and intramolecular topology, promoting membrane association consistent with its proposed neurotoxic activity.

### ΔCR shifts the N-terminus toward a membrane-engaged state

A first, robust signature of the ΔCR form is the increased propensity of the extreme N-terminus to approach and persist at the membrane surface. Both the K23–membrane distance distributions and the cumulative contact-time analysis show that ΔCR samples shorter distances and longer membrane-associated states than WT. This behavior is consistent with the long-standing view that the flexible N-terminus can act as a “toxic effector” whose activity is normally restrained by intramolecular regulation from the C-terminal domain, and that ΔCR mutants escape this regulation and become constitutively active in cells, where they induce spontaneous ionic currents and neurodegeneration ^7,8,12^. Importantly, in our simulations this increased membrane engagement occurs without requiring unfolding of the C-terminal core, reinforcing a mechanism in which toxicity is driven by altered domain-domain communication and membrane interactions rather than by global destabilization. Our end-to-end analysis further supports this framework. While both WT and ΔCR ensembles are more compact than a neutral-chain WLC reference, WT displays a smaller average end-to-end distance and a lower effective persistence length than ΔCR, consistent with a more compact conformational ensemble. The WT N-terminus more often samples configurations that are less permissive for membrane association, whereas ΔCR shifts the ensemble toward more extended states that can more readily reach the bilayer. Taken together, the distance and contact-time metrics as well as the WLC description converge on the interpretation that ΔCR biases the N-terminus toward conformations that enhance its membrane proximity and association.

### ΔCR rewires intra–N-terminal contact networks

The intra-domain contact analysis shows that, in WT, the highest-occupancy N-ter/N-ter contacts are strongly enriched in interactions involving CR residues, with frequent couplings between residues below 100 and the 108– 124 segment, and recurrent contacts linking the 50–60 region to residues just above 105. In ΔCR, all contacts involving residues 105–125 are necessarily lost, and the most persistent intra-N-ter interactions redistribute within the domain, with prominent couplings between the 30–40 and 80–90 regions and a dense cluster centered on the Gly-rich segment. Peak occupancies are reduced, consistently with a less “locked-in” intra-N-ter organization. Mechanistically, these patterns suggest that CR residues in WT do not merely contribute local contacts, but act as a hub that stabilizes couplings within the N-terminus. Removing this hub forces the N-terminus to reorganize around alternative contact motifs, yielding an ensemble with altered internal topology and slightly reduced maximal contact persistence. This rewiring is expected to change which parts of the N-terminus are exposed, which are sequestered, and how easily the chain can extend toward external interaction partners such as the membrane.

### WT and ΔCR adopt distinct N-ter/C-ter interaction patterns

Beyond shifting the N-terminal ensemble, the ΔCR deletion appears to also reshape the interplay between the disordered N-terminus and the globular domain. In WT, the N-ter/C-ter interface is not defined by a single rigid pattern; rather, it is supported by a set of contacts that persist for substantial portions of the dynamics and converge on a preferred interaction mode. This type of distributed inter-domain coupling provides the basis for intramolecular regulation: it can restrain the N-terminus near the globular domain surface, reducing the likelihood that the N-terminus resides at the membrane.

### ΔCR disrupts this regulatory architecture

The deletion removes an internal organizational element of the N-terminus and, in doing so, shifts the inter-domain interaction propensity toward alternative interaction surfaces on the C-terminal domain. Functionally, this remodeling is likely to reduce the propensity of the C-terminal domain to sequester the N-terminus near the protein surface, thereby increasing the population of conformations in which the N-terminus is more available for membrane association. Notably, ΔCR samples that partially recover WT-like inter-domain coupling also display reduced N-terminal membrane engagement, suggesting that membrane association and inter-domain restraint are dynamically coupled and compete on the same conformational landscape.

### The deletion of CR reprograms PrP conformational landscape

Overall, these results converge on a unified mechanistic picture in which the central region acts as a critical intramolecular hub that governs the balance between N-terminal sequestration and membrane engagement. Its deletion does not weaken the system in a simple manner, but instead rewires the conformational landscape at multiple levels, loosening intra–N-terminal organization, redistributing inter-domain contacts, and ultimately shifting the ensemble toward states in which the N-terminus is more exposed, extended, and membrane-active. In this framework, ΔCR PrP emerges not as a destabilized protein, but as a reprogrammed one, in which altered long-range communication between domains favors a membrane-engaged topology. This provides a structurally grounded explanation for how central region deletions can convert PrP into a constitutively active, neurotoxic species without requiring global unfolding, and highlights the central region as a key regulator of PrP functional and pathological states.

## Data Availability

All the data that support the findings of this study are available within the manuscript, supplementary files, or from the corresponding authors upon reasonable request.

## Code Availability

Codes developed in this study are available from the corresponding authors upon reasonable request.

## Acknowledgements

We thank all the members of the Dulbecco Telethon Laboratory of Prions & Amyloids at Department CIBIO, University of Trento (www.cibio.unitn.it/95/dulbecco-telethon-laboratory-ofprions-and-amyloids) for the fruitful discussion of the results presented in this manuscript. We thank Jesús R. Requena for the access to supercomputing facilities Finis Terrae III (CESGA, Spain).

This work was supported by grants from Fondazione Telethon (GGP20043, GMR24T2072), the American CJD Foundation and the Italian Association for Prion Encephalopathies (AIENP) to E.B.

## Author Contribution

Conceived and designed the computational analyses: all authors

Conceived and designed the experimental analyses: n/a

Performed the computational analyses: MR

Performed the experimental analyses: n/a

Analysed the data: MR, PF, EB

Contributed reagents/materials/analysis: n/a

Wrote the paper: all authors

Edited the paper: all authors

## Supplementary Figures

**Figure S1.**
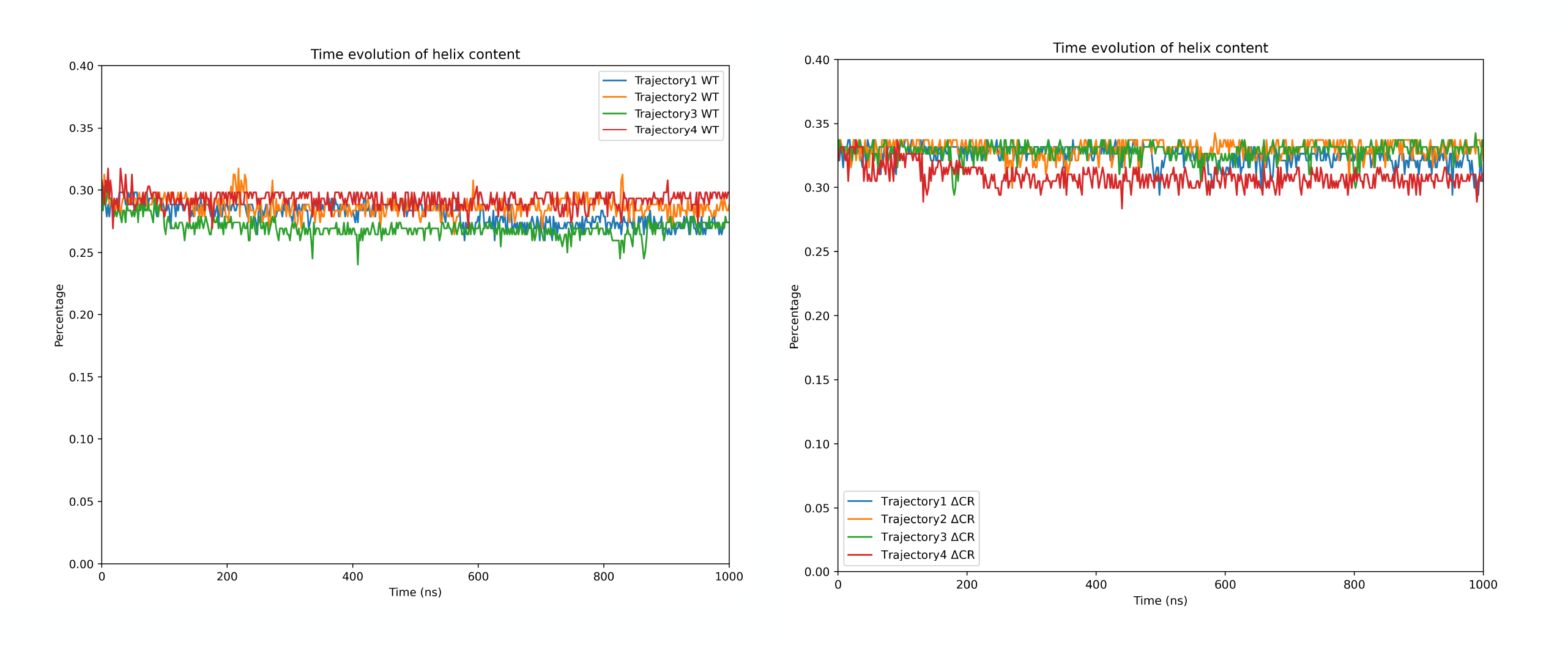
Time evolution of the total content of helix structure during the simulations. The percentage of helix content is expressed as a value between 0 and 1, and represents the number of residues in helix conformation over the total number of residues. The ΔCR PrP system has a higher percentage of helix content with respect to WT PrP due to the shorter chain.

**Figure S2.**
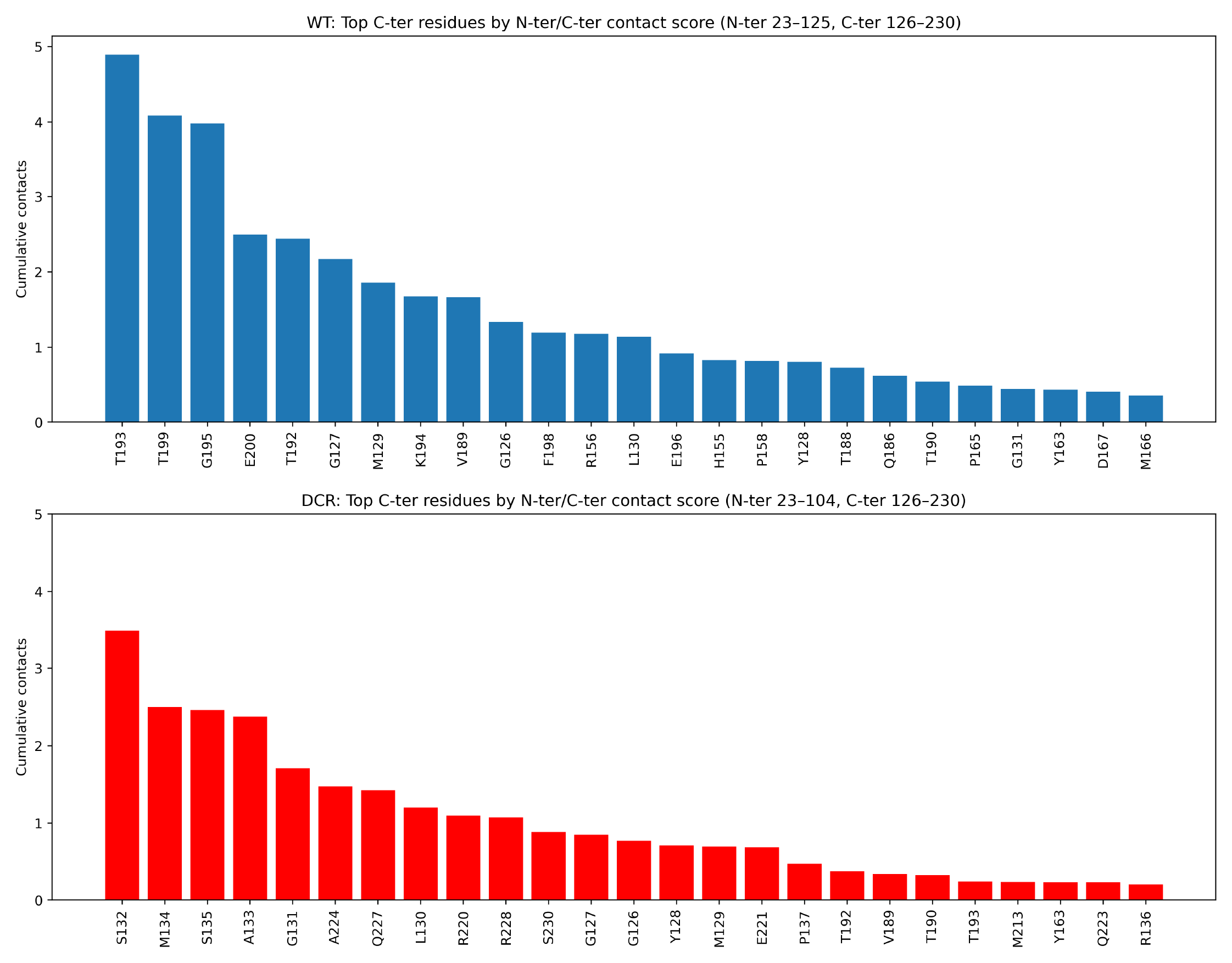
Top C-terminus residues involved in N/C contacts in WT or ΔCR PrP. Bars report the C-terminal contact score for each residue, defined as the sum of mean contact fractions over all N-terminal partners (i.e., cumulative N/C contact occupancy per C-terminal residue). WT PrP is computed over N-terminus 23–125 and C-terminus 126–230, whereas ΔCR PrP is computed over N-terminus 23–104 (Δ105–125) and the same C-terminus 126–230.

**Figure S3.**
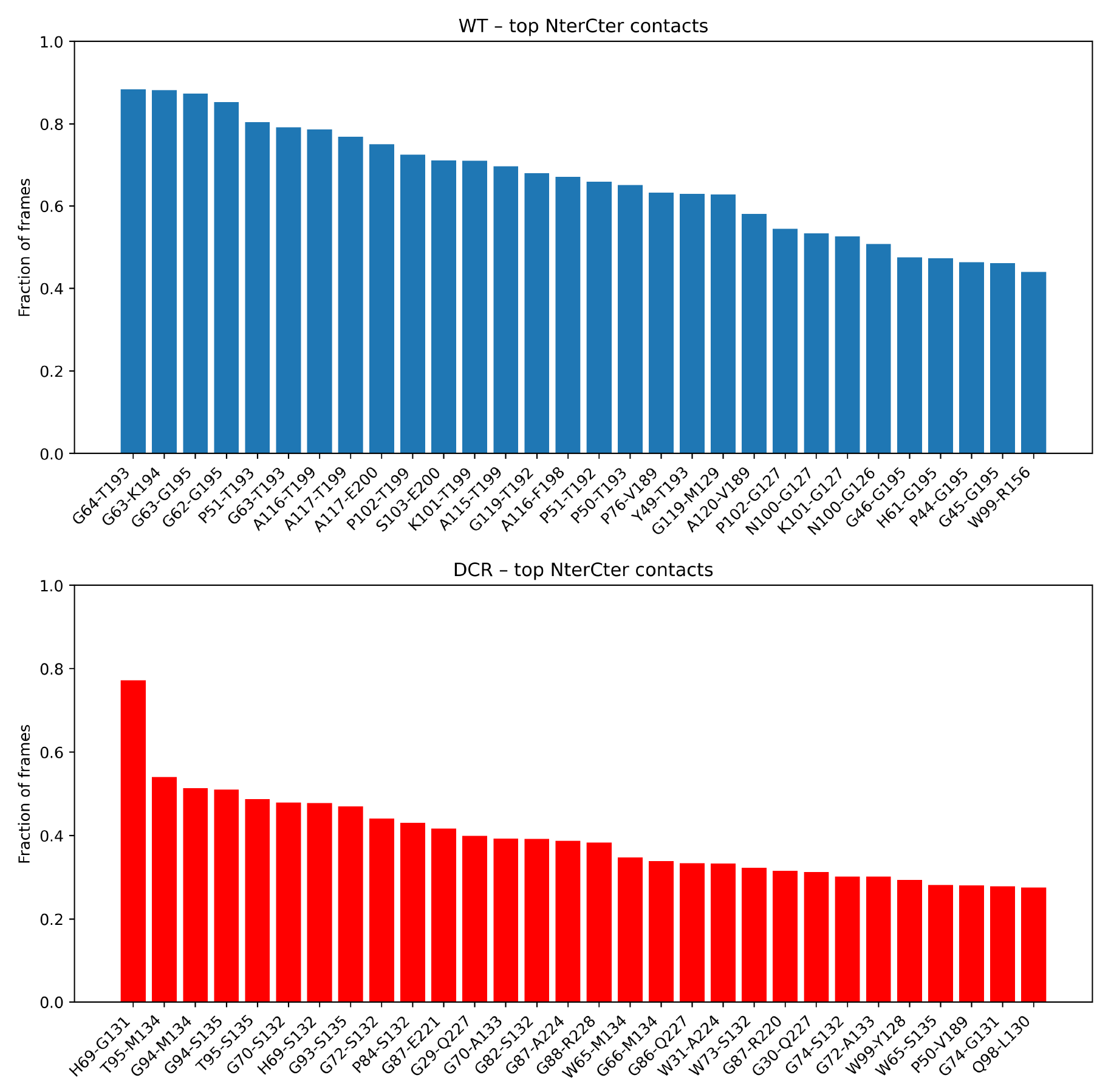
Most persistent N/C residue pairs in WT or ΔCR PrP molecules. Bars show the mean contact fraction (calculated as the fraction of trajectory frames where residues are in contact) for the top-ranked intra-domain residue pairs. WT PrP pairs predominantly involve the C-terminal hotspot around 192–200, whereas in ΔCR PrP the strongest contacts shift toward the 131–135 region and a secondary region near 220–228, with overall reduced peak occupancies.

**Figure S4.**
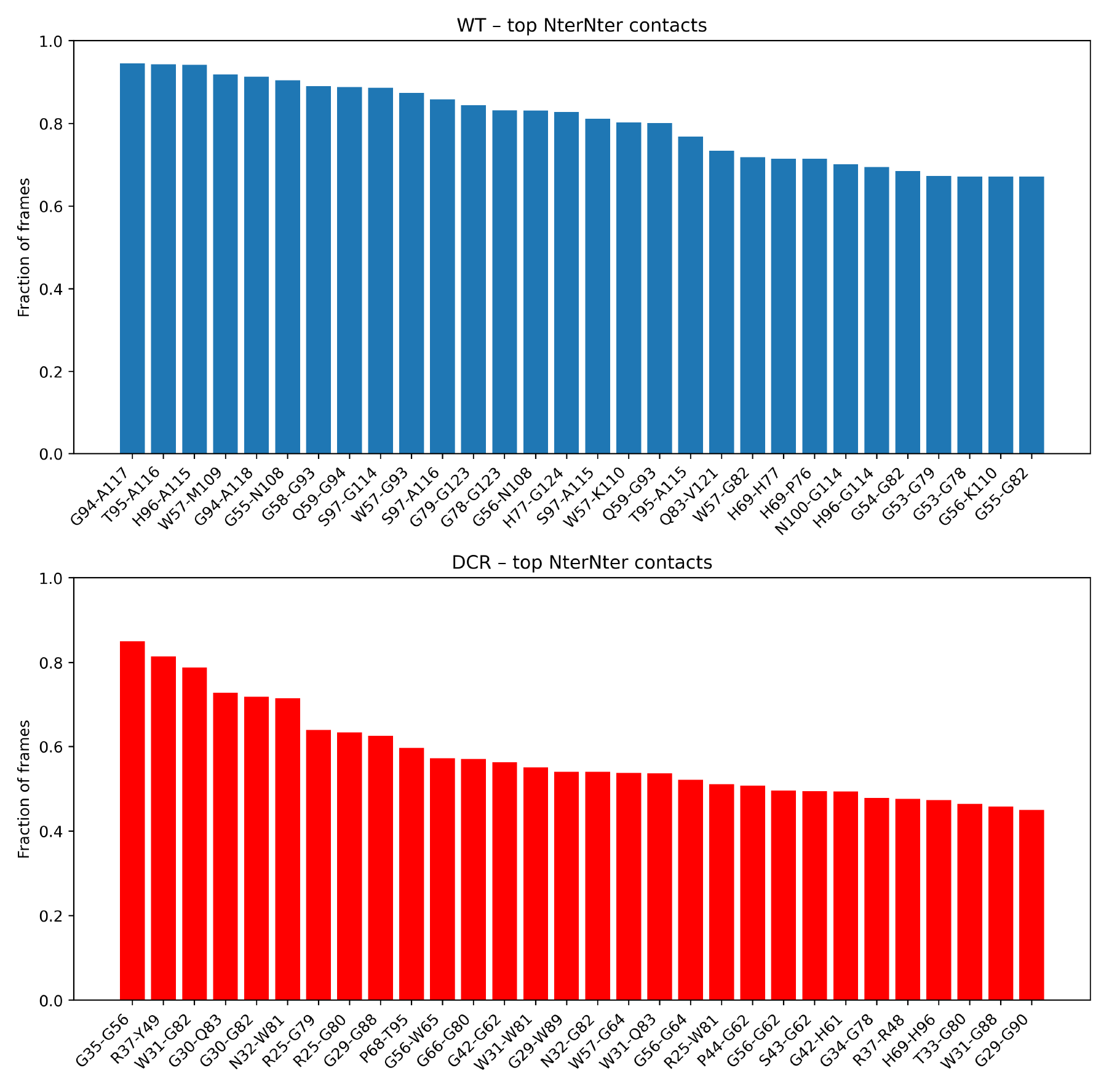
Top N/N contacts in WT or ΔCR PrP molecules. Bars report the fraction of frames in contact for the highest-ranking intra–N-terminal residue pairs. In WT PrP (N-ter 23–125), many top contacts involve residues within CR, whereas in ΔCR (N-ter 23–104; Δ105–125) the top contacts are between the 30–40 and 80–90 segments and around the mid-N-terminal region (∼56–66).

**Supplementary V1**. The video illustrates a single MD trajectory of membrane-bound full-length WT (left panel) and ΔCR (right panel) PrP showing distinct conformational behaviors. While WT PrP adopts a more compact ensemble stabilized by interactions between the N-terminal and globular C-terminal domains, the ΔCR mutant displays extended conformations, loss of key intramolecular contacts, and increased N-terminal association with the membrane surface.

## References

1. Prusiner, S. B. Prions. Proceedings of the National Academy of Sciences 95, 13363–13383 (1998).

2. Biasini, E., Turnbaugh, J. A., Unterberger, U. & Harris, D. A. Prion protein at the crossroads of physiology and disease. Trends Neurosci. 35, 92–103 (2012).

3. Linden, R. et al. Physiology of the Prion Protein. Physiol. Rev. 88, 673–728 (2008).

4. Reimann, R. R. et al. Differential Toxicity of Antibodies to the Prion Protein. PLoS Pathog. 12, e1005401 (2016).

5. Sonati, T. et al. The toxicity of antiprion antibodies is mediated by the flexible tail of the prion protein. Nature 501, 102–106 (2013).

6. Massignan, T., Biasini, E. & Harris, D. A. A Drug-Based Cellular Assay (DBCA) for studying cytotoxic and cytoprotective activities of the prion protein: A practical guide. Methods 53, 214–219 (2011).

7. Solomon, I. H., Biasini, E. & Harris, D. A. Ion channels induced by the prion protein: mediators of neurotoxicity. Prion 6, 40–45 (2012).

8. Solomon, I. H., Huettner, J. E. & Harris, D. A. Neurotoxic mutants of the prion protein induce spontaneous ionic currents in cultured cells. J Biol Chem 285, 26719–26726 (2010).

9. Solomon, I. H. et al. An N-terminal polybasic domain and cell surface localization are required for mutant prion protein toxicity. J Biol Chem 286, 14724–14736 (2011).

10. Li, A. et al. Neonatal lethality in transgenic mice expressing prion protein with a deletion of residues 105-125. Embo J 26, 548–558 (2007).

11. Baumann, F. et al. Lethal recessive myelin toxicity of prion protein lacking its central domain. Embo J 26, 538–547 (2007).

12. Solomon, I. H., Schepker, J. A. & Harris, D. A. Prion neurotoxicity: insights from prion protein mutants. Curr Issues Mol Biol 12, 51–62 (2009).

13. Iraci, N., Stincardini, C., Barreca, M. L. & Biasini, E. Decoding the function of the N-terminal tail of the cellular prion protein to inspire novel therapeutic avenues for neurodegenerative diseases. Virus Res. 207, 62–68 (2015).

14. Varadi, M. et al. AlphaFold protein structure database in 2024: Providing structure coverage for over 214 million protein sequences. Nucleic Acids Res. 52, D368–D375 (2023).

15. Lee, J. et al. CHARMM-GUI Input Generator for NAMD, GROMACS, AMBER, OpenMM, and CHARMM/OpenMM Simulations Using the CHARMM36 Additive Force Field. J. Chem. Theory Comput. 12, 405–413 (2016).

16. Huang, J. et al. CHARMM36m: An improved force field for folded and intrinsically disordered proteins. Nat. Methods 14, 71–73 (2017).

17. Lindahl, Abraham Hess & Spoel, van der. GROMACS 2021 Manual. https://doi.org/10.5281/ZENODO.4457591 doi:10.5281/ZENODO.4457591.

18. Evans, D. J. & Holian, B. L. The Nose–Hoover thermostat. J. Chem. Phys. 83, 4069–4074 (1985).

19. Parrinello M., R. A. Polymorphic transitions in single crystals: A new molecular dynamics method. J. Appl. Phys. 52, 7182–7190 (1981).

20. Darden, T., York, D. & Pedersen, L. Particle mesh Ewald: An N·log(N) method for Ewald sums in large systems. J. Chem. Phys. 98, 10089–10092 (1993).

21. Hess B., B. H. B. H. J. C. F. J. G. E. M. LINCS: A linear constraint solver for molecular simulations. J. Comput. Chem. 18, 1463–1472 (1997).

22. Young, P. Physics 115/242 Leapfrog method and other ‘symplectic’ algorithms for integrating Newton’s laws of motion.

23. Doi, M. . & Edwards, S. F. . The theory of polymer dynamics. 391 (20041986).

24. Mühle, S., Zhou, M., Ghosh, A. & Enderlein, J. Loop formation and translational diffusion of intrinsically disordered proteins. Phys. Rev. E 100, 052405 (2019).

25. Humphrey, W., Dalke, A. & Schulten, K. VMD: Visual molecular dynamics. J. Mol. Graph. 14, 33–38 (1996).

26. Meng, E. C. et al. UCSF ChimeraX: Tools for structure building and analysis. Protein Sci. 32, (2023).

27. Martinez, J. et al. PrP charge structure encodes interdomain interactions. Scientific Reports 2015 5:1 5, 13623–(2015).

